# Whole genome sequencing of orofacial cleft trios from the Gabriella Miller Kids First Pediatric Research Consortium identifies a new locus on chromosome 21

**DOI:** 10.1101/743526

**Authors:** Nandita Mukhopadhyay, Madison Bishop, Michael Mortillo, Pankaj Chopra, Jacqueline B. Hetmanski, Margaret A. Taub, Lina M. Moreno, Luz Consuelo Valencia-Ramirez, Claudia Restrepo, George L. Wehby, Jacqueline T. Hecht, Frederic Deleyiannis, Azeez Butali, Seth M. Weinberg, Terri H. Beaty, Jeffrey C. Murray, Elizabeth J. Leslie, Eleanor Feingold, Mary L. Marazita

## Abstract

Orofacial clefts (OFCs) are one of the most common birth defects worldwide and create a significant health burden. The majority of OFCs are non-syndromic, and the genetic component has been only partially determined. Here, we analyze whole genome sequence (WGS) data for association with risk of OFCs in European and Colombian families selected from a multicenter family-based OFC study. Part of the Gabriella Miller Kids First Pediatric Research Program, this is the first large-scale WGS study of OFC in parent-offspring trios. WGS provides deeper and more specific genetic data than currently available using imputation on single nucleotide polymorphic (SNP) marker panels. Here, association analysis of genome-wide single nucleotide variants (SNV) and short insertions and deletions (indels) identified a new locus on chromosome 21 in Colombian families, within a region known to be expressed during craniofacial development. This study reinforces the ancestry differences seen in the genetic etiology of OFCs, and the need for larger samples when for studying OFCs and other birth defects in admixed populations.

## Introduction

Orofacial clefts, primarily cleft lip (CL) and cleft palate (CP) are among the most common birth defects in all populations worldwide with differences in birth prevalence by ancestry (1, 2). Surgical treatment along with ongoing orthodontia, speech and other therapies, are very successful in ameliorating the physical health effects of OFC, but there is still a significant social, emotional and financial burden for individuals with OFC, their families, and society (3, 4). Furthermore, there are disparities in access to such therapies for OFCs (5), similar to other malformations with complex medical and surgical needs. Some studies have suggested a reduced quality of life for individuals with OFCs (6), while other studies have identified higher risk to certain types of cancers (7–9). Thus, it is critical to identify etiologic factors leading to OFCs to improve diagnostics, treatments, and outcomes.

The causal genes for most syndromic forms of OFCs are now known, and listed within OMIM (https://www.ncbi.nlm.nih.gov/omim, search criterion=(cleft lip cleft palate syndrome) AND “omim snp”[Filter]), but the majority of OFC cases - including about 70% of CL with or without CP (CL/P) and 50% of CP alone - are considered non-syndromic, i.e. they occur as isolated anomalies with no other apparent cognitive or structural abnormalities (1). The causal genes for non-syndromic OFCs are still largely undiscovered. To date, there have been 52 genome-wide associations reported and replicated between non-syndromic CL/P and genetic markers (NHGRI-EBI Catalog of published genome studies) (10), but as for most other complex human traits (11–13), very few putative functional variants for non-syndromic OFCs have been identified from genome-wide association studies (GWASs)(14). In particular, the high heritability for OFC, estimated at 90% by a twin study in a Danish sample (15) cannot be explained by all identified common variants significantly associated with OFC, sometimes referred to as the “missing heritability” problem (16). Additional approaches will be necessary to expand our understanding of genetic variation in nonsyndromic OFCs and whole genome sequencing (WGS) holds the promise of teasing out the so-called missing heritability from GWASs of OFC and other complex traits (17).

An important new approach has been implemented by the Gabriella Miller Kids First Pediatric Research Consortium (https://commonfund.nih.gov/kidsfirst/overview). Kids First was established in 2015 to address gaps in our understanding of the genetic etiologies of structural birth defects and pediatric cancers by providing WGS of case-parent trios with these major pediatric conditions. Addressing both of these areas (structural birth defects and pediatric cancers) in Kids First was partially motivated by the observation that children with birth defects such as OFCs are at a higher risk of also developing some cancers, and their family members also have elevated risk (7, 8), suggesting there may be shared genetic pathways underlying cancer and birth defects. The KidsFirst study consists of 952 case-parent trios (i.e. affected probands and their parents) from multiple OFC studies, of which, 415 are of European descent, 275 Latino, 125 Asian and 137 African. The current study summarizes initial findings on common variants, i.e. single nucleotide polymorphic (SNP) markers and small insertions/deletions from WGS of a sample of 315 trios European descent, as well as a sample of 265 trios of Latin American ancestry from Colombia, all with offspring affected with cleft lip with or without cleft palate (CL/P).

## Results

### Genome-wide association of SNPs and indels

Genome-wide associations using allelic and genotypic transmission disequilibrium test (TDT) were run separately in 315 **European** and 265 **Colombian** trios and then in the **Combined** set of all 580 trios on bi-allelic single nucleotide polymorphic (SNP) markers and indels with minor allele frequency (MAF) greater than 10% (see Methods for discussion of the MAF cutoff). A comparison of the p-values between allelic TDT (aTDT) and genotypic TDT (gTDT) showed high concordance (see section “<u>Comparison between aTDT and gTDT</u>” and supplementary figure S1 Figure 1), therefore, only the aTDT results are discussed in the following sections. P-values calculated using the exact binomial distribution from McNemar’s test are reported for the aTDT.

**Figure 1.**
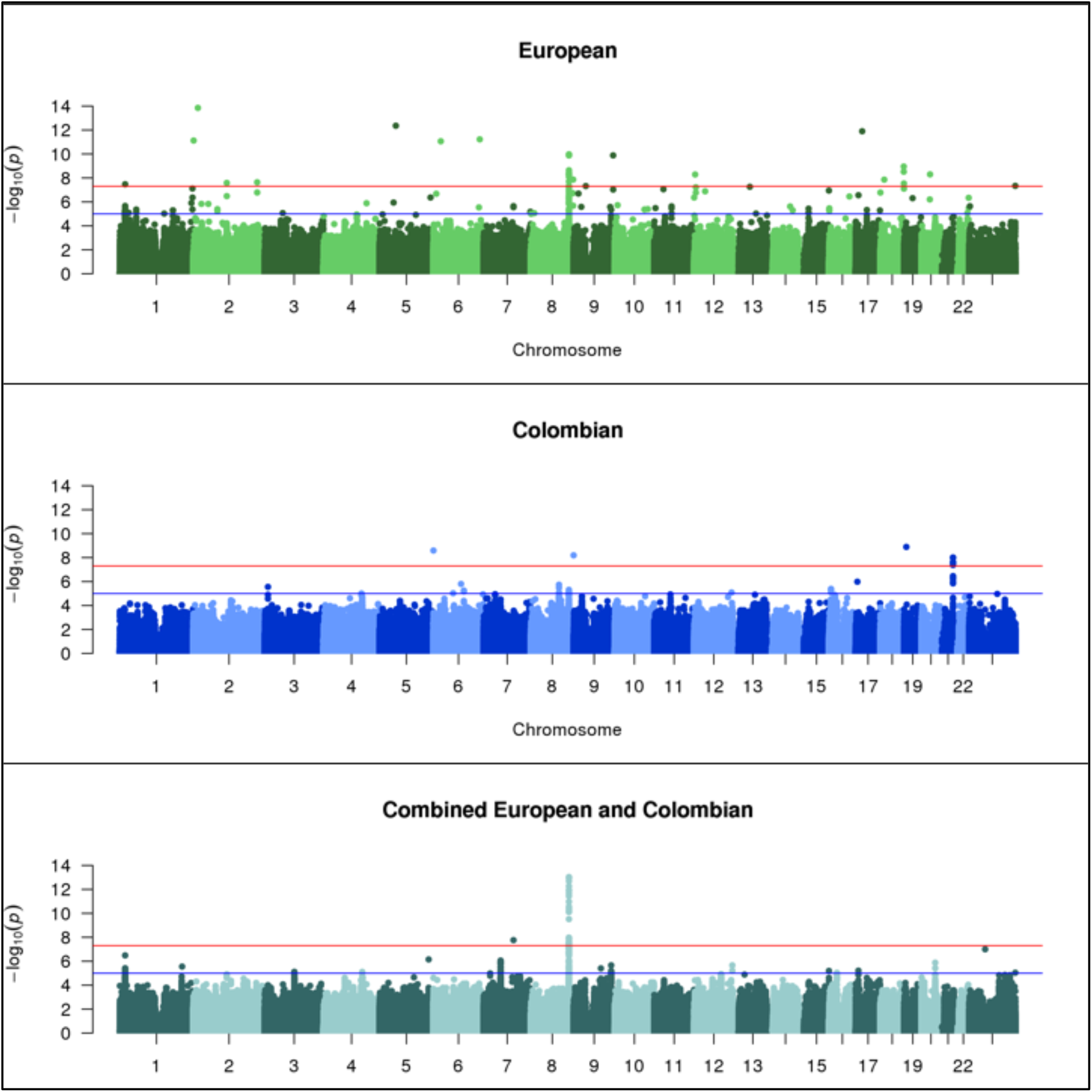
Manhattan plots of **European (315 trios)**, **Colombian (265 trios)** and **Combined (580 trios)** allelic TDTs

Tables 1 and 2 show the most significant results in the **European** (Table 1) and **Colombian** trios (Table 2). Several SNPs gave genome-wide significant association p-values in the stratified aTDT analysis of **European** (Table 1 and Figure 1 top panel) and **Colombian** trios (Table 2 and Figure 1 middle panel), and a single SNP achieved genome-wide significance in the **Combined** sample (Figure 1 bottom panel). In the **European** sample, 17 significant associations are observed across multiple chromosomes (Table 1). In the **Colombian** sample, four significant associations are observed for markers on chromosomes 6, 8, 19 and 21. After close examination of the genome-wide significant associations in the **European** and **Colombian** trios, the one strongly supported new result was a region on chromosome 21q22.3, discussed below. In the **Combined** aTDT, a single genome-wide significant association (p = 9.35E-14, OR = 2.13, 95% CI = [1.74–2.62], SNP rs72728755) was observed in the 8q24.21 chromosomal region. Many of the other associations showed properties that reduced our confidence in their reliability, which included (1) no additional variants yielding either significant or suggestive p-values close to the lead SNP, (2) the lead SNP was located in a highly repetitive region, or (3) the lead SNPs showed substantial differences in MAF across European or Latino samples in gnomAD (18). Therefore, we concluded that these might not be reliable signals. Note that the first criterion alone was not sufficient to make us deem a result unreliable, as the 10% MAF cutoff may have been responsible for single-SNP association peaks.

**Table 1.**
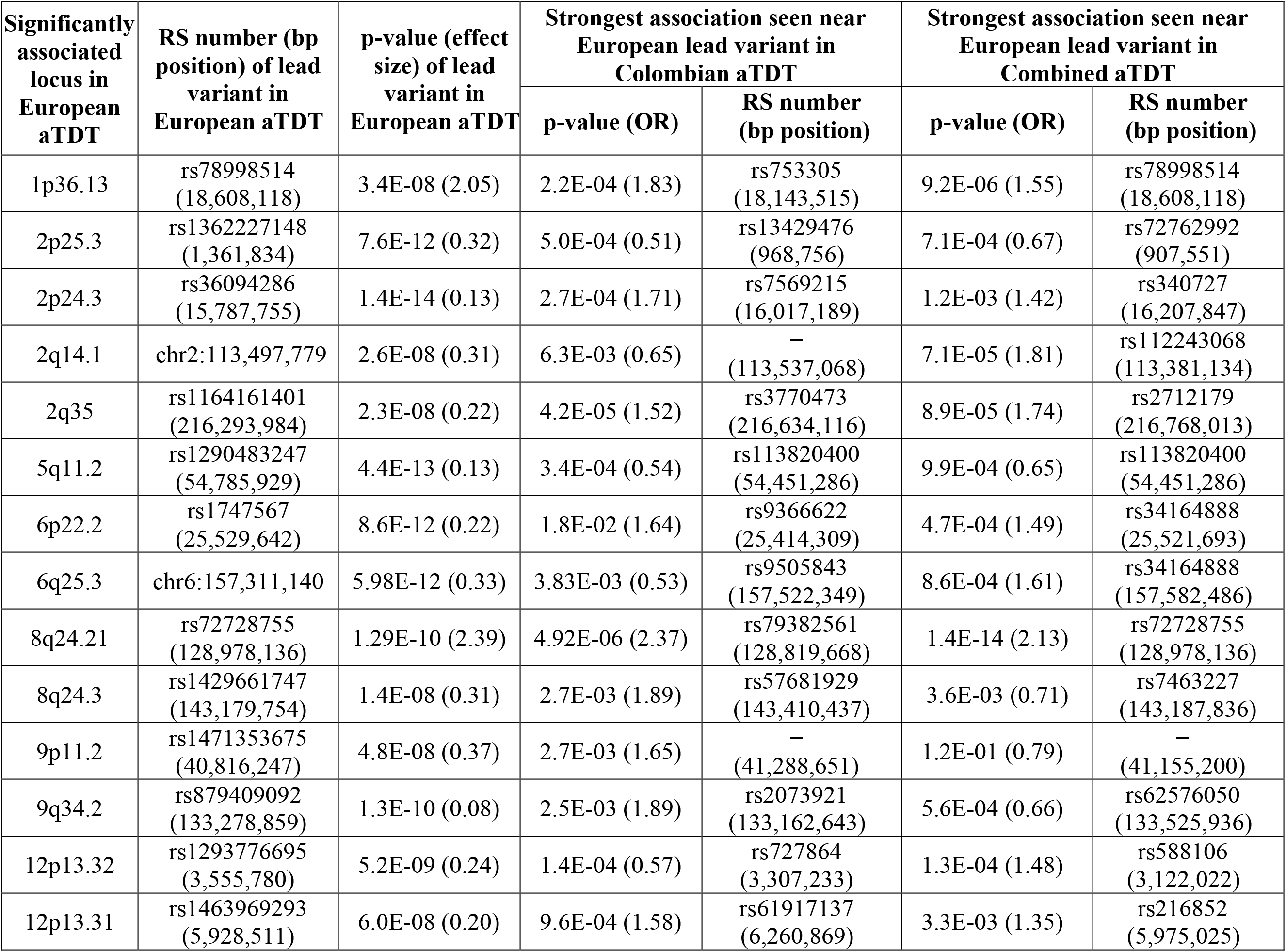
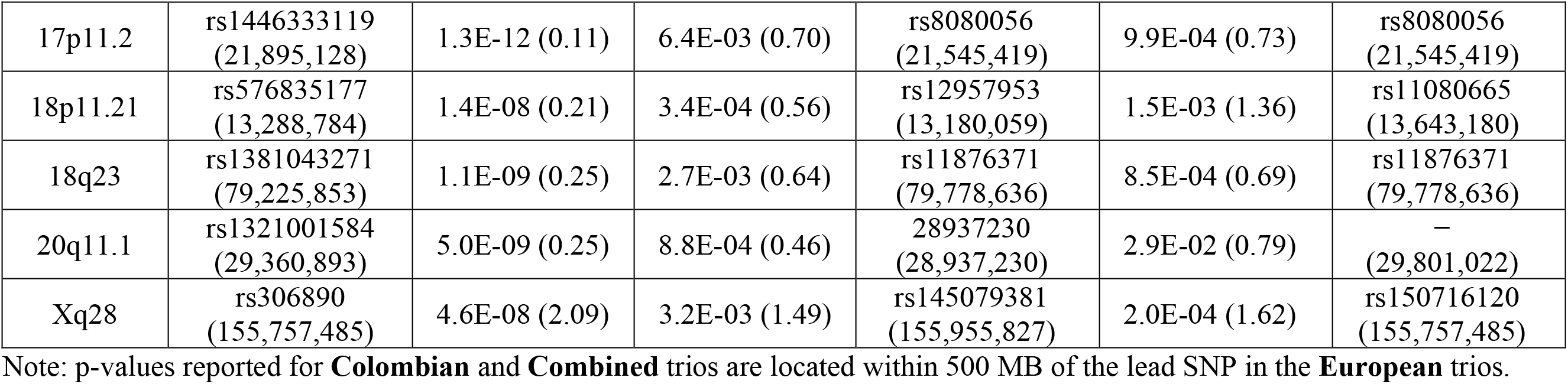
Significant associations in **European** (315 trios) compared with **Colombian** (265 trios) and **Combined** (580 trios).

**Table 2.**
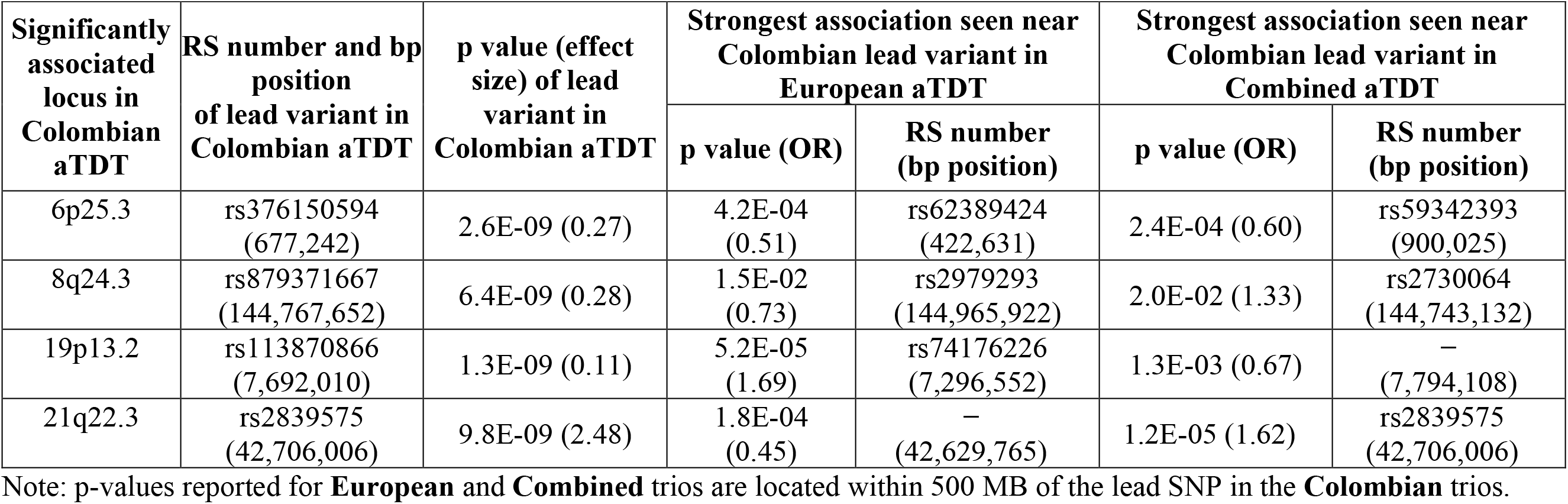
Significant associations in **Colombian** (265 trios) compared with **European** (315 trios) and **Combined** (580 trios).

| Significantly associated locus in Colombian aTDT | RS number and bp position of lead variant in Colombian aTDT | p value (effect size) of lead variant in Colombian aTDT | Strongest association seen near Colombian lead variant in European aTDT |  | Strongest association seen near Colombian lead variant in Combined aTDT |  |
| --- | --- | --- | --- | --- | --- | --- |
|  |  |  | p value (OR) | RS number (bp position) | p value (OR) | RS number (bp position) |
| 6p25.3 | rs376150594<br>(677,242) | 2.6E-09 (0.27) | 4.2E-04 (0.51) | rs62389424<br>(422,631) | 2.4E-04 (0.60) | rs59342393<br>(900,025) |
| 8q24.3 | rs879371667<br>(144,767,652) | 6.4E-09 (0.28) | 1.5E-02 (0.73) | rs2979293<br>(144,965,922) | 2.0E-02 (1.33) | rs2730064<br>(144,743,132) |
| 19p13.2 | rs113870866<br>(7,692,010) | 1.3E-09 (0.11) | 5.2E-05 (1.69) | rs74176226<br>(7,296,552) | 1.3E-03 (0.67) | –<br>(7,794,108) |
| 21q22.3 | rs2839575<br>(42,706,006) | 9.8E-09 (2.48) | 1.8E-04 (0.45) | –<br>(42,629,765) | 1.2E-05 (1.62) | rs2839575<br>(42,706,006) |

Note: p-values reported for **European** and **Combined** trios are located within 500 MB of the lead SNP in the **Colombian** trios.

### Comparison between allelic TDTs of European and Colombian trios

A qualitative comparison of the **European** and **Colombian** aTDT results showed few commonalities between the two analyses of common SNPs. Except for the peaks at the 8q24.3 chromosomal region, all other genome-wide significant regions in the **European** trios were neither significant nor suggestive in the **Colombian** trios, and vice versa. The lack of new signals from the **Combined** trios supports this observation. For the purposes of comparison, Table 1 lists all **European** peaks and contains the smallest association p-values with their corresponding estimated odds ratios (OR) observed in the **Colombian** and **Combined** aTDTs within 500 KB on either side of each **European** peak SNP (Table 1 columns 4-7). Since allele frequencies for specific SNPs may differ between the two samples, this provides a region-level view of replication across the samples. Similarly, Table 2 lists the **Colombian** peaks, along with the minimum association p-values and corresponding odds ratios observed in the **European** and **Combined** aTDTs within 500 KB on either side of each **Colombian** peak. As seen in Tables 1 and 2, **European** and **Colombian** trios differ considerably with respect to the genomic regions that show significant association to CL/P.

### Previously reported OFC risk loci

Two of the genome-wide significant associations observed in this study, 1p36.13 and 8q24.21, have been previously reported as associated with risk to OFCs by our group and others (19–21). The 1p36.13 peak is located 23kb upstream of the transcription start site of the *PAX7* gene. These associations were significant only in our **European** trios, consistent with previous studies suggesting a stronger association in participants of European ancestry compared to other racial/ethnic groups (22).

The 8q24.21 region has been consistently implicated in nearly all previous OFC studies especially among samples of European ancestry. The lead SNP among **Europeans** (rs55658222) is in strong linkage disequilibrium (LD) with another SNP rs987525 in the HapMap European sample. The rs987525 SNP was found to be the lead SNP in this region in several previous GWASs. This SNP also showed modest evidence of association and linkage in the **Colombian** trios (p-value 8.609e-06, odds ratio=1.984, CI= [1.46–2.69]). In the **European** trios, a suggestive association was observed for an indel located at 9,295,770 bp on chromosome 17, approximately 52kb centromeric to the *NTN1* gene (p=2.77e-07, odds ratio=0.29, CI=[0.18–0.48]). None of the other previously reported OFC variants reached even a suggestive level of significance (suggestive threshold p<1.0e-05) in our WGS study, which is not unexpected given the smaller sample size of this WGS study compared to published GWASs. Supplement S2 Table shows the most significant aTDT p-values within 500 KB of all previously reported OFC risk variants.

### Chromosome 21q22.3 association in the Colombian trios

We observed genome-wide significant associations in the **Colombian** trios within a 30kb interval on chromosome 21q22.3 (Figure 2, top panel). In this sample, the common variants had relatively large estimated odds ratios ranging from 2.33 to 2.48, i.e. approximately two-fold increases in the transmission of the risk alleles from parents to the proband offspring. The smallest p-value was observed at rs2839575 (p=9.75e-09, odds ratio=2.48, 95% CI = [1.81–3.45]).

**Figure 2.**
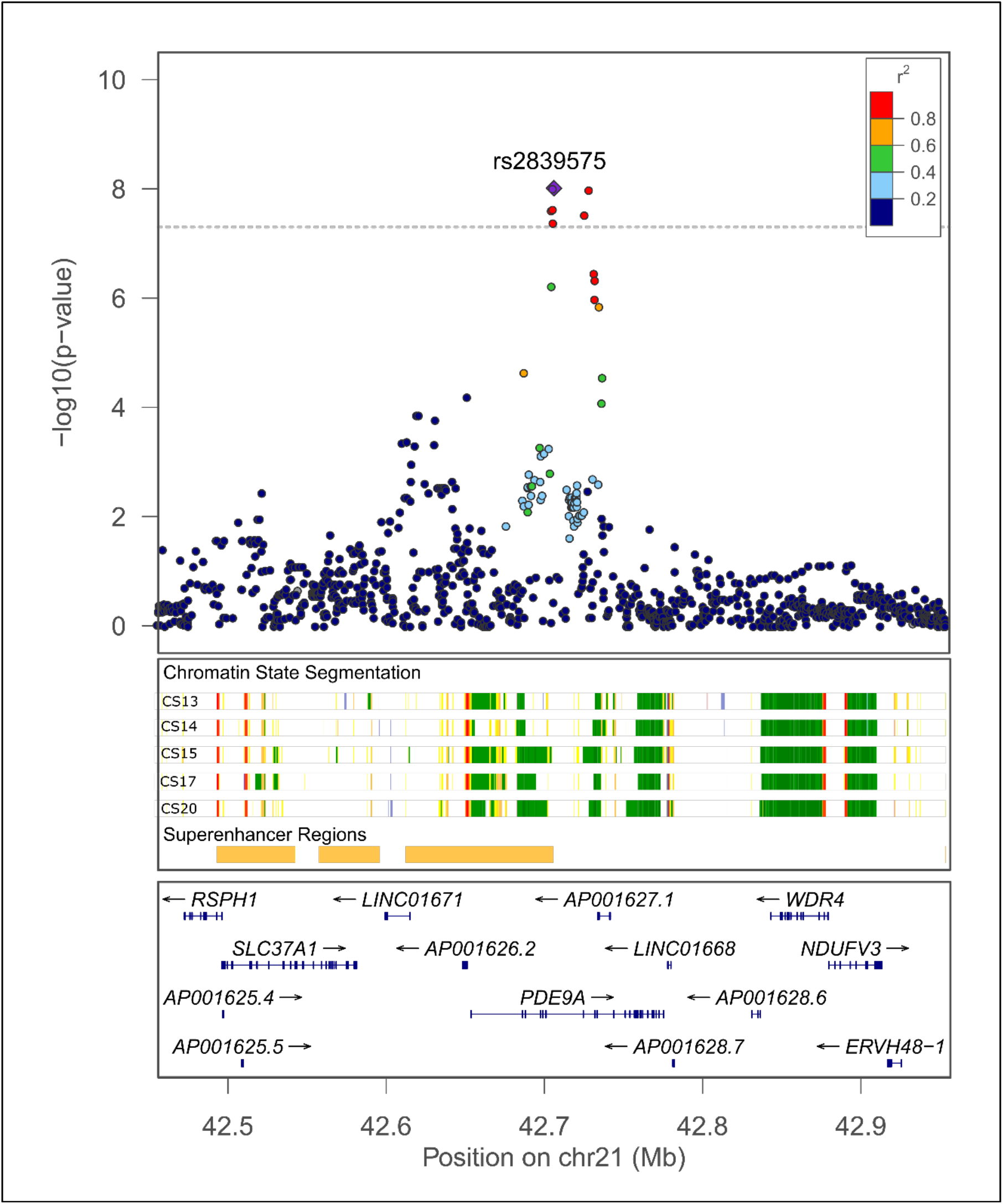
Log_10_(p) for SNPs and indels in chromosome 21q22.3 peak region.

GWAS of a Latino sample from a previous study, the POFC Multiethnic study (24), reported suggestive association at this genomic region (see Figure 1 in (23)). That Latino sample included diverse Hispanic groups from the US, Guatemala, Argentina, and Colombia, and all of the current WGS **Colombia** trios were part of the POFC Multiethnic study. However, the POFC Multiethnic study had 129 additional Colombian trios. In that study, the GWASs of Asian and European samples did not show association in this region, and nor did the combined GWAS of all the POFC Multiethnic study samples. The fact that the current WGS case-parent trio study yielded a genome-wide significant association with a smaller sample size suggests this association might be unique to **Colombians**. We explored the validity and implications of this observation through a number of analyses, as described below.

We first examined the aTDT p-values for our **Colombian** WGS trios using their SNP array data from the POFC Multiethnic study. The p-values in this region were nearly identical to those observed in our WGS association, confirming the association we observed here was not an artifact of sequencing.

We next investigated whether population substructure within the Colombian parents could have caused the observed association in the WGS data by examining the ancestry principal components (PCs) as well as results of quantitative association between PCA eigenvalues to variants within the peak region (see Methods for details). PCA showed no evidence of population substructure (supplementary figure S1 Figure 2A), and no association was observed between the eigenvalues and variants in the chromosome 21q22.3 region (supplementary figure S1 Figure 2B). A positive association between eigenvalues and variants would have indicated that the observed association with CL/P is in reality due to population substructure, therefore, this association did not appear to be an artifact of population admixture.

We verified that this region does not show evidence of association in other Latin Americans, by reanalyzing imputed genotype data from the previously published POFC Multiethnic GWAS study (23). P-values from the aTDT of independent trios and the corresponding odds ratios at rs2839575 in each Latino subpopulation were considerably different from those observed in the Colombian subjects (Figure 3A, forest plot). In fact, in contrast to the Colombians, none of the other Latino populations (except Colombia) showed a significant association at rs2839575. Moreover, the combined set of non-Colombian Latinos resulted in much weaker associations across a 1 MB region flanking SNP rs2839575 as well as for this SNP itself. The odds ratios at the rs2839575 variant showed an opposite (although non-significant) effect in the non-Colombian Hispanics as compared to Colombians (Figure 3B, regional p-value plot and supplementary table S3 Table). We concluded from the stratified aTDT results that this SNP influences OFC risk only in Colombians.

**Figure 3.**
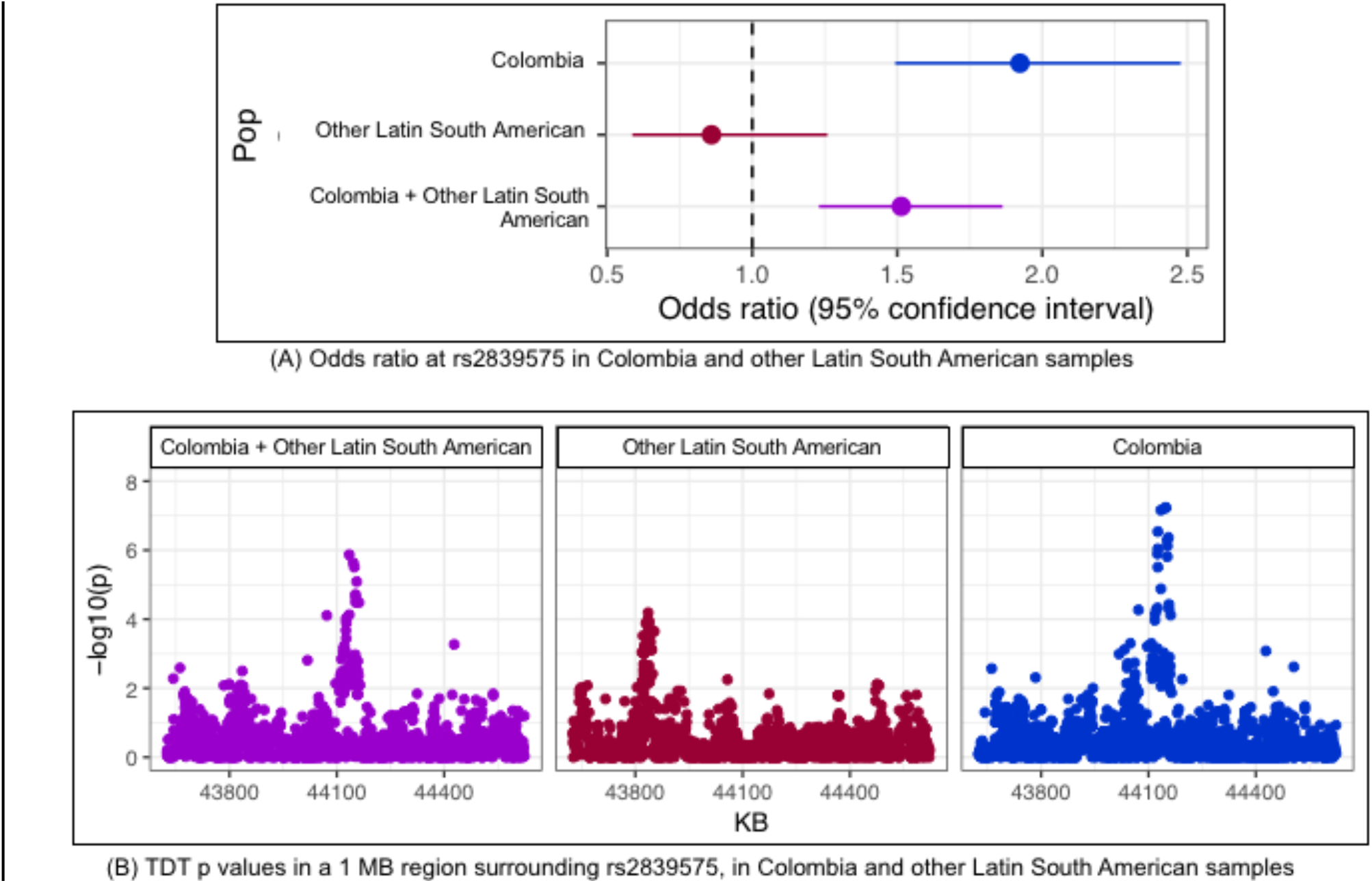
Estimated odds ratios (with 95%CI) and −log10(p-values) from the aTDT in Colombia and other Latino samples

We therefore investigated the possibility of ancestry differences between our Colombian sample and the other Latino populations. Ancestry principal components calculated from the POFC Multiethnic SNP genotype data (unrelated individuals only), showed Colombians to be ancestrally diverse from the other Latino populations (supplementary figure S1 Figure 3).

Given that the 21q22.3 association is observed only in the Colombian sample and that the ancestry of Colombians is different from the other Latin American samples, we checked whether the absence of an association signal in the other Latin American samples merely reflects differences in MAF rather than differences in true effects of risk alleles. That is, it is possible that a causal variant exists in all populations but has a considerably higher frequency (or is in LD with a variant of higher frequency) in the Colombians. Given the population history of Colombians, causal OFC variants are may have arisen from one particular ancestral group, and such variants may be more frequent (and therefore more informative) among Colombians. The origin of African ancestry of Colombians is different from that of the other Latino populations (24). We therefore looked at the frequencies of the Colombian risk alleles across different populations. For this analysis, we again turned to genotyped and imputed SNP genotypes from the POFC Multiethnic study. The MAFs of the 30 most significantly associated SNPs within the 21q22.3 peak region in Colombian trios were compared to 15 populations defined by country of recruitment from the POFC Multiethnic study. None of these 30 SNPs had higher MAF among Colombians compared to other Latino populations (supplementary figure S1 Figure 4). Moreover, the 15 most significant SNPs in this peak region had higher allele frequencies in all other population groups (European, African, and Asian) compared to Colombians or other Latinos. Thus, there was no conclusive evidence that population-specific variants contributed to the association signal seen in this study. However, several of these variants had estimated odds ratios between 1.1 and 1.5 in Asian, Europeans, or Africans, suggesting these variants in this region may also increase risk for OFCs in other populations, but at a reduced level.

Finally, we tested for effects of rare variants within the **Colombian** trios using burden and collapsing tests because we observed a number of low-frequency and rare variants with large odds ratios in this region (see Methods for rare variant testing procedure). Common variants with the strongest associations were all intronic variants within the *PDE9A* gene; however, all had moderate odds ratios around 2.0. In this region, there were 37 SNPs with minor allele frequencies near or below 1% in the **Colombian** trios and estimated OR > 5 (supplementary table S4 Table), including mainly intronic and a few intergenic SNVs (28 intronic, 8 intergenic). The exception was one non-synonymous SNV, rs138007679 in the *RSPH1* gene (aTDT odds ratio 8, 95% CI = [1.001–63.96]), which produces an amino acid change (A>C, leucine to tryptophan according to ClinVar). Alone, this variant does not clearly implicate *RSPH1* over other genes in the region, so we performed a rare variant TDT on all non-synonymous variants within the 13 genes falling in this region. None of the individual genes achieved the nominal significance (supplementary table S5 Table), so this result remained inconclusive. We also carried out rare variant TDTs of intronic and intergenic variants with similar results, finding only nominally significant associations attributable to intergenic, low frequency variants (MAFs ranging between 0.5% and 1%).

In the absence of any clearly pathogenic variant or gene based on combined effects of rare variants, we examined regulatory elements and protein-protein interaction pathways in this region with respect to craniofacial development. All associated variants below a suggestive level of significance (p<1.0e-05) were located within the *PDE9A* gene, which does not have any known role in controlling risk to OFCs. However, the *PDE9A* gene overlaps a super-enhancer region for craniofacial development identified from histone profiling in early human craniofacial development (25). Multiple genes in the region, including *PDE9A*, appear to be actively transcribed during human craniofacial development (Figure 2). Another gene of interest is *UBASH3A*, located ~220kb centromeric to this peak signal. The UBASH3A protein was previously shown to physically associate with *SPRY2* via a yeast two-hybrid assay (33). SPRY2 has been reported by GWASs of OFC and shown required for palatogenesis in mice (26); whether UBASH3A is also expressed in craniofacial structures has not yet been determined.

## Discussion

This study is the first large scale WGS study of OFCs, one of the most common birth defects worldwide, using a case-parent trio design. We conducted association analyses of common variants from WGS in two samples of case-parent trios, one of European ancestry and the other of Latin American ancestry from Colombia. We replicated two known OFC loci and identified a promising new region on chromosome 21 in the Colombian sample. A combined association analysis of these two samples together clearly shows that OFC risk loci differ by ethnicity. The 8q24 locus has been repeatedly shown to be associated with risk of OFCs in both case-control and case-parent trio samples from a range of ethnicities such as Europeans and Latin Americans, with some evidence from Asians (27). Here, we found slight differences in the larger 8q24 region between Europeans and Latin Americans but there appears to be a shared risk locus at 8q24.21, consistent with Colombians having a strong influence from European ancestry. *IRF6*, a gene that has been linked to OFCs in samples of Asian and Latino ancestry was not detected in our Colombian trios, possibly due to the small sample size.

We observed evidence of linkage and association to a previously unreported region on chromosome 21 spanning the *PDE9A* gene only in the Colombian sample. We verified that this locus is unique to Colombians, by running separate aTDTs in Colombian and non-Colombian Latino trios using imputed genotype data from the previous POFC Multiethnic GWAS study (23). We examined whether the apparent risk alleles have ancestral origins from non-Latino populations and noted that the estimated effect sizes were slightly elevated in Asian, European and African populations although never achieving genome-wide significance. However, larger or more phenotypically specific samples may be necessary to find conclusive statistical evidence. The significantly associated common variants in the chromosome 21q22.3 peak were mostly intronic or intergenic, with no obvious biological function. There were a number of rare variants with large aTDT odds ratios, including a non-synonymous SNP within the *RSPH1* gene, however, TDT of rare coding non-synonymous variants did not provide conclusive statistical evidence of association between genes in this region and CL/P. Although none of the genes in this region are known to contribute to the development of OFCs, they appear to be actively transcribed during human craniofacial development and should be examined further in follow up studies.

## Research Design and Methods

### Study design

Two samples of case-parent trios were analyzed for the current family-based association study, one of European descent recruited from sites around the United States, Argentina, Turkey, Hungary and Madrid, and a second of trios from Medellin, Colombia. The two samples are referred to as **European** and **Colombian** respectively, in this study. Recruitment of participants and phenotypic assessments were done at regional treatment centers for orofacial clefts after review and approval by the site-specific IRBs.

This study included case-parent trios consisting of affected offspring and their parents (see Table 3). Most of the **European** parents and all **Colombian** parents are unaffected for CL/P (see breakdown of trios in Table 3). All trios had offspring with a cleft lip or a cleft lip plus cleft palate, and had not been diagnosed with any recognized genetic syndrome. The affection status was defined as cleft lip with or without cleft palate (CL/P) for all analyses here because the Colombian sample did not have the breakdown between cleft lip alone (CL) versus cleft lip with cleft palate (CLP). Table 3 shows the counts of GMKF trios sequenced for the present study, by their country of origin.

**Table 3.** Counts of CL/P trios by recruitment site and cleft type (add?)

| Sample | Total Trios | Trios with no affected parents | Trios with 1 affected parent | Trios with 2 affected parents |
| --- | --- | --- | --- | --- |
| European | 315 | 280 | 32 | 3 |
| Site: USA | 209 | 185 | 21 | 3 |
| Hungary | 56 | 51 | 5 |  |
| Madrid | 30 | 26 | 4 |  |
| Argentina | 1 | 1 |  |  |
| Turkey | 19 | 17 | 2 |  |
| **Colombian** | 265 | 265 |  |  |
| **Total** | 580 | 545 | 32 | 3 |

### Genetic data

Whole genome sequencing of the European sample was carried out at the McDonnell Genome Institute (MGI), Washington University School of Medicine in St. Louis, while sequencing of the Colombian sample was conducted at the Broad Institute, both with an average of 30X coverage. Variant calling on the European trios was performed using pipelines at MGI, and aligned to the GRCh37/hg19 genome assembly. The European sample’s genotypes were realigned and recalled by the GMKF’s Data Resource Center at Children’s Hospital of Philadelphia to match the Colombian sample, which was aligned to hg38 and called using GATK pipelines (28–30) at the Broad Institute (https://software.broadinstitute.org/gatk/best-practices/workflow). The alignment and joint genotyping workflow used to harmonize these two samples of case-parent trios was developed using GATK Best Practice recommendations, with the goal of being functionally equivalent with other current large genomic research efforts. Briefly, the harmonization pipeline first converted the mapped alignments within each sample to unmapped alignments, then re-ran the GATK genotyping workflow, namely base quality score recalibration (BQSR), simultaneous calling of SNPs and indels using single sample variant calling (HaplotypeCaller), multiple sample joint variant calling, and finally refinement and filtering of called variants. Data processing and storage of harmonized results was done on the Cavatica platform within an Amazon Web Services (AWS) environment. The GMKF Data Resource Center (DRC) was responsible for tracking, final checking, and release of the variant calls via its portal. The released variant data contained genotypes called at 35,600,754 single nucleotide variants (SNVs) and 4,320,146 indels mapped to the hg38 reference sequence. Details of the harmonization process are provided in the supplement, S1 section “Kids First DRC Genomics Harmonization Pipeline Description”.

### Assessment of sample data quality and data cleaning

Each sample of trios (**European, Colombian**) was separately analyzed for genotyping inconsistencies, at an individual level, as well as on a trio basis. **Genotype quality**: Genotypes with either unacceptable read depth (minimum depth 10 reads for autosomes; minimum 5 reads for X chromosomes in males), or genotyping quality (minimum GQ 20; minimum GQ=10 for X chromosome variants in males) were first set to unknown. **Sample quality:** Each individual’s set of variant calls was checked for excess heterozygosity (> 3 standard deviations from mean heterozygote/homozygote ratio), deviant transition to transversion ratios (Ts/Tv > 3 standard deviations from mean Ts/Tv across samples), low genotyping rates (below 90%), and for inconsistency between the average homozygosity on the X-chromosome and the individual’s reported sex. Each trio was assessed for Mendelian error rates and deviation from the expected degree of relatedness between each set of parents and offspring. Genomes flagged for sex or relationship issues were compared with SNP array genotypes from the POFC Multiethnic study (23) to resolve sample swaps or misclassification of sex, where possible (some trios from our study were not part of POFC Multiethnic study). A trio was excluded if it failed more than one of these data quality tests, and if recovery was not possible after comparison with the SNP array genotype data.

After QC procedures, the final dataset consisted of 315 complete European trios and 265 complete Colombian trios. Biallelic variants including SNPs and short indels (indels range between 1-10,000 BP in length) with a genotyping rate of at least 90% were included in our analyses. A total of 5,374,579 variants were analyzed in the **European** trios, and 4,905,638 in **Colombian** trios. Of these, 4,220,712 variants were analyzed for the **Combined** trios.

### Genome-wide wide association testing of SNPs and indels

Genome-wide association was conducted using two versions (allelic and genotypic) of the transmission disequilibrium test (TDT), for each polymorphic variant. The PLINK software (31, 32) was used to run the standard genome-wide allelic TDT (aTDT), which does not consider the parents’ cleft status. We also ran genotypic TDT (gTDT) (33) on the trios, and compared the association p-values to those from the aTDT. Effect sizes are not directly comparable between the two methods. The aTDT compares the transmission of a target allele to the affected child from heterozygous parents (34), and is based on McNemar’s chi-square statistic. Because only heterozygous parents can contribute to this statistic, statistical power is greatly influenced by minor allele frequency (MAF) and one population may have considerably more or less power at any given SNP when MAF varies across populations. The gTDT compares the observed genotype in the child to “pseudo-controls” representing other genotypes possible from the parental mating type. Schwender et al. (35, 36) demonstrated an efficient method for computing this gTDT statistic. Because either TDT represents a test of strict Mendelian inheritance of the marker (despite sampling case-parent trios through the affected proband), this test is robust to spurious associations arising from population stratification and can provide greater power for rare phenotypes (37). The null hypothesis of either TDT is the complete absence of either linkage between the marker and an unobserved causal gene or linkage disequilibrium (LD) between the marker and an unobserved causal gene. Rejection of this composite null hypothesis implies the presence of both linkage and LD. The TDT is most appropriate for our study, given our participants originate from diverse populations, and the Colombians in particular are known to reflect varying degrees of admixture of African, Hispanic, Native American and European genes.

Three genome-wide TDT analyses were run: separately in European and Colombian trios and then in all trios combined. Significance p-values for the allelic TDT statistic were calculated using the exact binomial distribution. Although the TDT statistic is robust to population substructure, an overall TDT analysis can mask subgroup specific results, thus principal component analysis (PCA) was run on the parents separately for each sample (**European**, **Colombian**) and the normalized eigenvalues examined for evidence of sub-groups within each sample. For PCs producing eigenvalues exceeding ± 5, we conducted genetic association assuming an additive model using the eigenvalues of each individual as quantitative traits. The PCA was conducted using the KING program (38). PLINK (31, 32) was used to run the quantitative association.

### Identification of significant associations

Due to our limited sample sizes, only SNPs with a minor allele frequency of at least 10% within each sample of trios were considered in these TDT analyses. The allelic TDT test relies on asymptotics, and can give inflated associations for lower MAF SNPs at this sample size when applied genome-wide. We subsequently examined lower-MAF SNPs in specific regions for fine-mapping purposes (see below). The genome-wide threshold for significant association was set at 5.0e-08, and the critical value for suggestive association was set at 1.0e-05.

### Fine mapping and rare-variant association in 21q region

A subset of the genome-wide significant associations (i.e. those not overlapping with previously reported OFC genes/regions) was selected for more in-depth investigation. All biallelic variants with a genotyping rate of 90% or greater, regardless of MAF, were investigated within each region of interest (defined as 1Mb centered on each lead variant). Each interval was annotated for possible roles in craniofacial development by literature searches of all genes contained within that interval, functional annotation of variants using multiple tools including Bystro (39), Variant Effect Predictor (40), and HaploReg (41). We also queried the UCSC genome browser’s gene-by-gene interaction track for known OFC genes/regions. This track identifies genes reported in protein-interaction databases and recognized biological pathways (42).

Rare variant (RV) association using the TDT framework was run only for regions containing SNPs showing significant evidence of linkage and association in the aTDT. For each association peak, we identified all genes located within 500 KB of the lead SNP, and selected non-synonymous RVs within the exons of these genes. Burden and collapsing methods were used, as our dataset is composed solely of case-parent trios, and these tests were applied to each gene separately, after phasing the observed genotype data of common SNPs. Beagle was used to calculate haplotypes (43) using all variants within a selected region. The RV-TDT software (44) was then run on phased haplotypes for exonic, non-synonymous SNVs in genes with a minimum of 4 variant sites. RV-TDT reports burden and combined multivariate and collapsing (CMC)-types of test statistics, as well as a weighted sum statistic. The observed MAFs within European and Colombian parents were used to calculate weights for each RV, where SNVs with smaller MAFs receiving higher weights. Some of the RV-TDT statistics use phased haplotypes to calculate empirical p-values by permuting the haplotypes of each parent. In addition to the exonic rare variants, we also selected intronic and intergenic variants and analyzed these using RV-TDT. Intronic and intergenic variants were divided up into subsets based on gene locations in this region, and analyzed using a procedure similar to the exonic, non-synonymous SNPs.

## Supporting information

Supplementary figures and variant calling procedure

Comparison with known OFC loci

Regional plots of chromosome 21q22.3

Rare SNPs with odds ratio 5.0 and greater

Rare variant TDT results

SNPs showing suggestive association in European trios

SNPs showing suggestive associations in Colombian trios

SNPs showing suggestive association in all trios

## Acknowledgements

These studies are part of the Gabriella Miller Kids First Pediatric Research Program consortium (Kids First), supported by the Common Fund of the Office of the Director of the National Institutes of Health (www.commonfund.nih.gov/KidsFirst). Washington University’s McDonell Genome Institute was awarded an administrative supplement (3U54HG003079-12S1) to sequence structural birth defect samples including the European descent Orofacial Cleft samples for the current study funded through Kids First (X01-HL132363). Further, the Broad Institute Sequencing Center was awarded a grant (U24-HD090743) to sequence structural birth defect cohort samples including the Latin American Orofacial Cleft family samples for the current study funded through Kids First (X01-HL136465). The sequencing centers plus the Kids First Data Resource Center (kidsfirstdrc.org, supported by the NIH Common Fund through U2CHL138346) provided quality control analyses in support of this project.

The data analyzed and reported in this manuscript were accessed from dbGaP [www.ncbi.nlm.nih.gov/gap; **European trios**: dbGaP accession number phs001168.v2.p2; **Colombian trios**: dbGaP accession number phs001420.v1.p1] and from the Kids First Data Resource Center (kidsfirstdrc.org). Additional grants supported the assembling of the cohorts, collection of the phenotypic data and samples, and data analysis [R01-DE016148, R03-DE026469, R01-DE012472, U01-DD000295, R01-DE014581, R01-DE011931, R37-DE008559, R21-DE016930, and R01-DE015667, R03-DE027193, R00-DE025060].

Many thanks to the participating families and study teams worldwide without whom these studies would not be possible. Additional thanks to Andrew Lidral, Mauricio Arcos-Burgos, and Andrew Czeizel.

