## Supplementary figures and variant calling procedure for "Whole genome sequencing of orofacial cleft trios from the Gabriella Miller Kids First Pediatric Research Consortium identifies a new locus on chromosome 21"

Comparison between aTDT and gTDT

We compared the TDT outcomes from allelic TDT (aTDT) and genotypic TDT (gTDT) genomewide for the European sample (5,226,459 SNPs and indels) and Colombian sample (4,749,979 SNVs and indels) separately. While gTDT p-values are consistently smaller than the aTDT pvalues, the differences are very small, with p[aTDT]/p[gTDT] ratios ranging between 0.8 and 1.25. 855 SNVs in the Colombian sample and 672 SNVs in the European samples show more significant aTDT p-values than gTDT. In the Colombian sample, the aTDT p-values at 4 SNVs in the chromosome 21q22.3 locus are above the genomewide significance threshold whereas the gTDT p-values fall slightly below the significance threshold (Figure 1 below). Effect sizes reported by the two forms of TDT are not directly comparable.

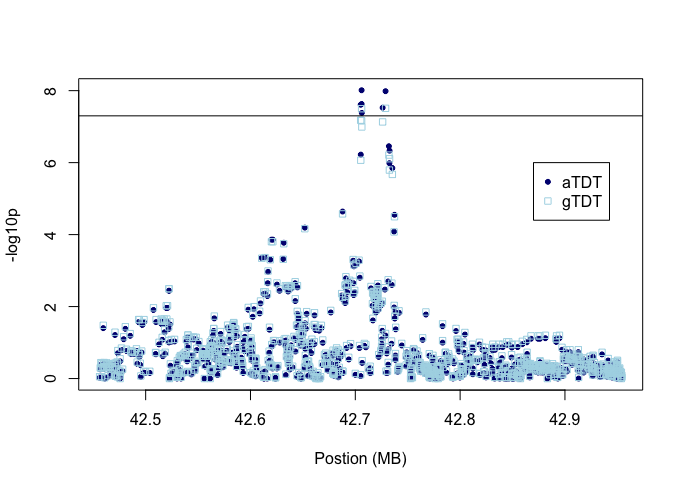

Figure 1. -log10 p-values from aTDT and gTDT analyses on chr21q22.3 SNPs

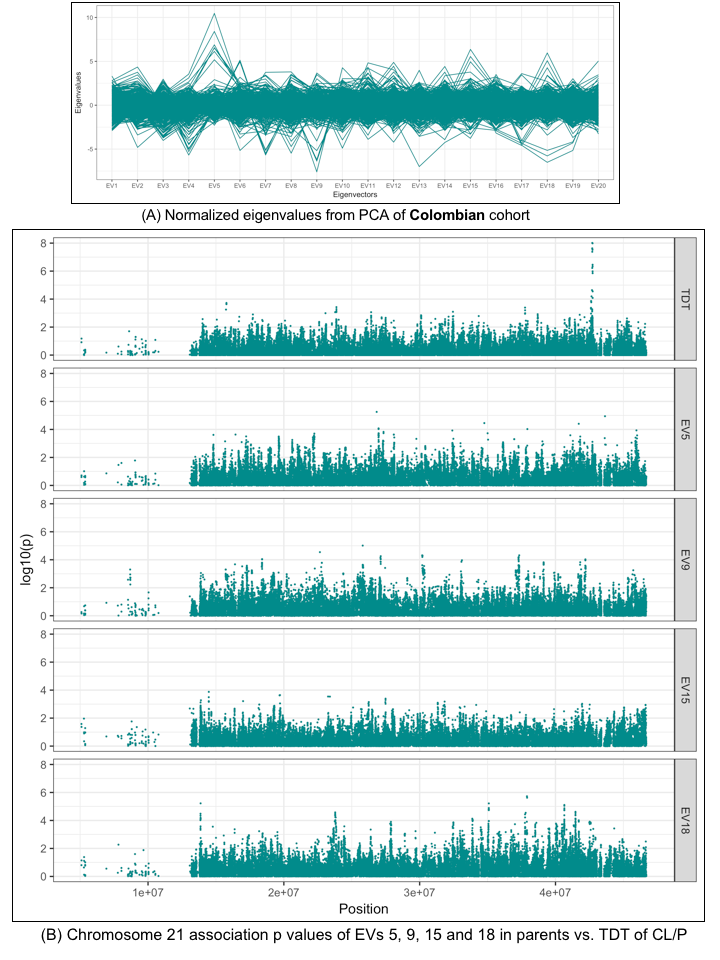
Figure 2. PCA and association of PCs in parents on chromosome 21

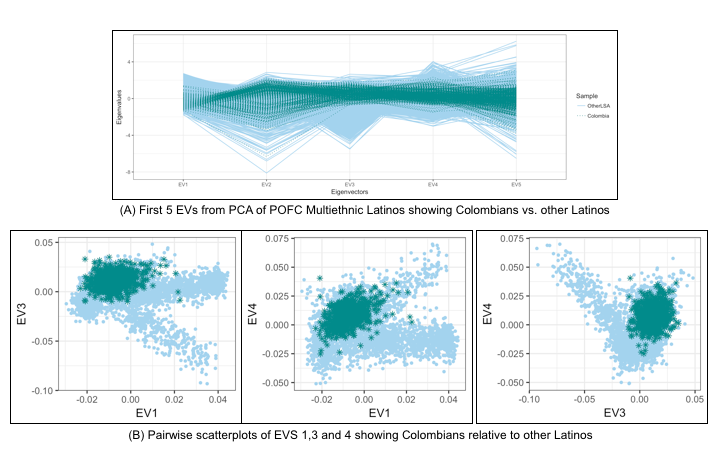

Figure 3. PCA Eigenvalues of Colombians vs. other Latinos in POFC Multiethnic study (Leslie et al. 2016)

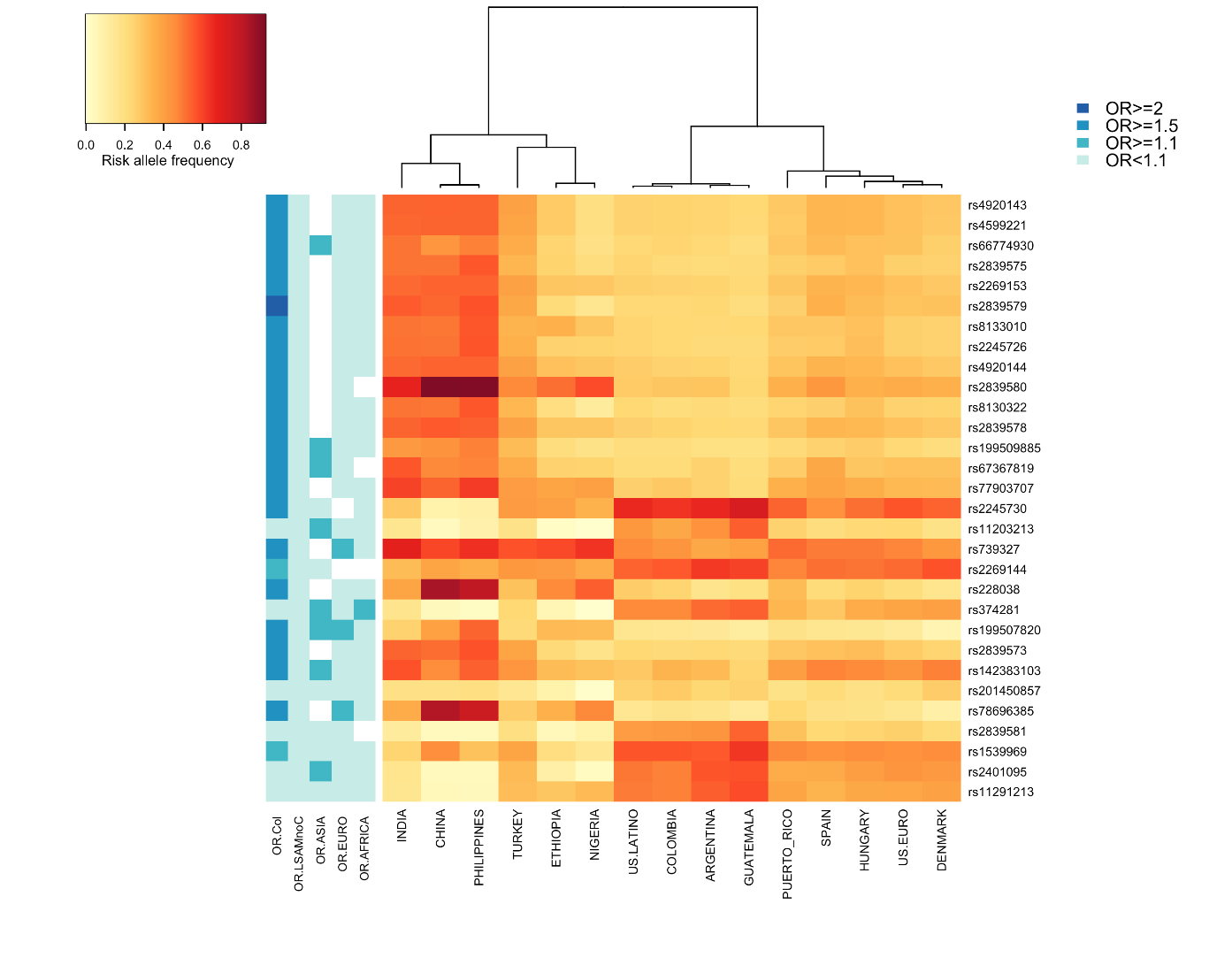

Figure 4. Risk allele frequency and odds ratio of the 30 most significantly associated SNPs from 21q22.3 in Multiethnic GWAS.

Figure 5. Q-Q plots of allelic TDT pvalues in European, Colombian and Combined samples

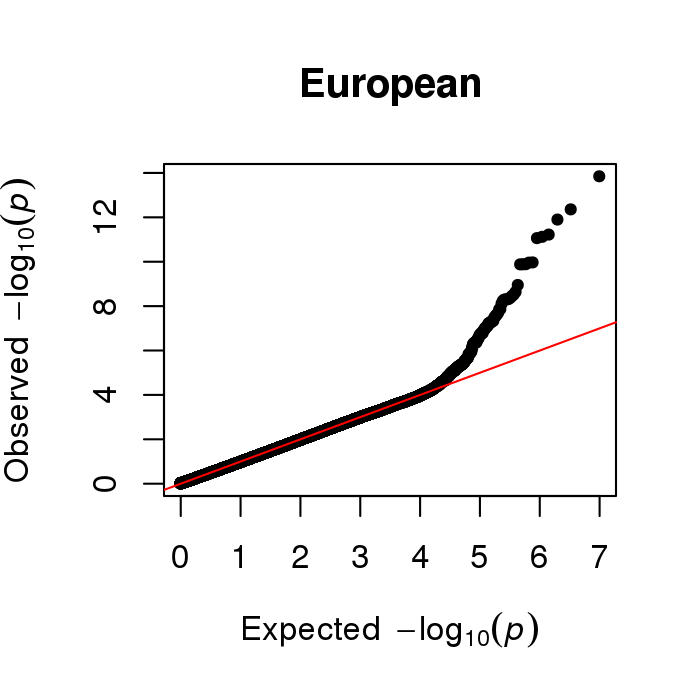

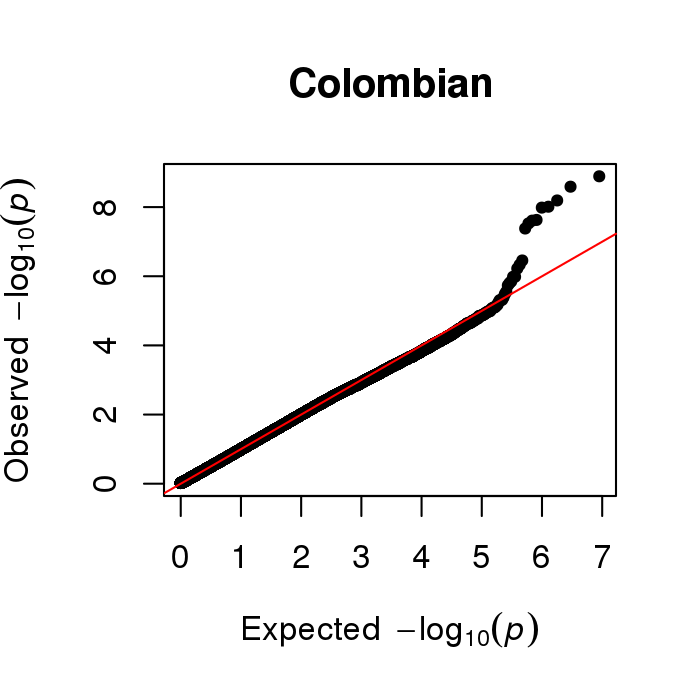

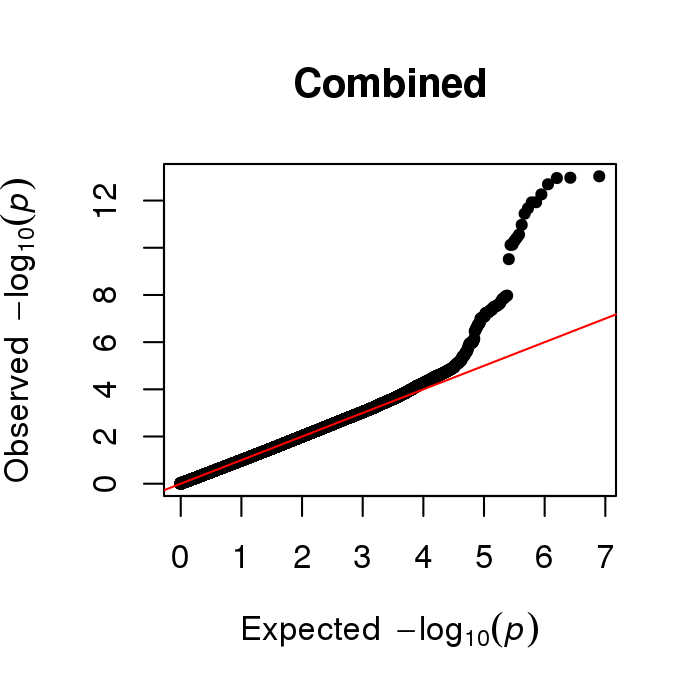

### Kids First DRC Genomics Harmonization Pipeline

The Kids First Data Resource Center (DRC) has developed and applied alignment and joint genotyping workflows following by the GATK Best Practice recommendations with the goal of being functionally equivalent with other current large genomic research efforts. The data processing is done via the Cavatica platform within an Amazon Web Services (AWS) environment. The harmonized results are stored in AWS and made searchable via the DRC Portal and further analyzable on Cavatica.

In more detail, the harmonization process started with an alignment workflow that takes input in the format of either BAM, CRAM or FASTQ or mixed, and converted them into uBAM (the unmapped BAM) by Picard RevertSam. After that, the uBAM were aligned, by readgroup, to human genome reference hg38, which included improved ALT contigs and HLA loci. Then Picard MarkDuplicates, SamSort and MergeBamAlignment were applied in a scatter execution fashion, where split chromosome intervals were run in parallel. Base Quality Score Recalibration (BQSR) was then applied. Lastly GATK4 HaplotypeCaller was applied to generate single sample gVCF along with the merged BAM converted into CRAM as final alignment outputs.

For the joint genotyping workflow, cohort-based gVCFs were imported as genomicsDB by GATK4 and passed down for GenotypeGVCFs execution. Variant Quality Score Recalibration (VQSR) was applied for SNP and InDel separately in a scatter fashion by calling intervals. A final VCF with a QC report were generated by GATK4 GatherVcfs and CollectVariantCallingMetrics. Finally, all the outputs were registered into the Kids First Data Service for tracking and checking of results and released to the DRC Portal and Cavatica, after final approval.

The following databases were applied during the BQSR and VQSR steps: for the SNPs, dbSNP138, HapMap, 1000 genomes and Omni; for the indels, dbSNP 138, Mills and axiomPoly. Kids First DRC pipelines are open source and made available to the public via GitHub:

Alignment workflow: https://github.com/kids-first/kf-alignment-workflow

Joint genotyping workflow: https://github.com/kids-first/kf-jointgenotyping-workflow
